# DRPnet - Automated Particle Picking in Cryo-Electron Micrographs using Deep Regression

**DOI:** 10.1101/616169

**Authors:** Nguyen P. Nguyen, Jacob Gotberg, Ilker Ersoy, Filiz Bunyak, Tommi White

## Abstract

Selection of individual protein particles in cryo-electron micrographs is an important step in single particle analysis. In this study, we developed a deep learning-based method to automatically detect particle centers from cryoEM micrographs. This is a challenging task because of the low signal-to-noise ratio of cryoEM micrographs and the size, shape, and grayscale-level variations in particles. We propose a double convolutional neural network (CNN) cascade for automated detection of particles in cryo-electron micrographs. Particles are detected by the first network, a fully convolutional regression network (FCRN), which maps the particle image to a continuous distance map that acts like a probability density function of particle centers. Particles identified by FCRN are further refined (or classified) to reduce false particle detections by the second CNN. This approach, entitled Deep Regression Picker Network or “DRPnet”, is simple but very effective in recognizing different grayscale patterns corresponding to 2D views of 3D particles. Our experiments showed that DRPnet’s first CNN pretrained with one dataset can be used to detect particles from a different datasets without retraining. The performance of this network can be further improved by re-training the network using specific particle datasets. The second network, a classification convolutional neural network, is used to refine detection results by identifying false detections. The proposed fully automated “deep regression” system, DRPnet, pretrained with TRPV1 (EMPIAR-10005) [1], and tested on *β*-galactosidase (EMPIAR-10017) [2] and *β*-galactosidase (EMPIAR-10061) [3], was then compared to RELION’s interactive particle picking. Preliminary experiments resulted in comparable or better particle picking performance with drastically reduced user interactions and improved processing time.

## 1 Introduction

Although high resolution 3D protein structure determination via single particle analysis (also known as single particle reconstruction) using cryo-electron microscopy (cryoEM) is becoming more widely used, it still remains a challenging technique because the resulting micrographs, especially those containing high resolution information, are noisy, usually with very low contrast intensity [4] [5]. In these cryoEM experiments, a purified, homogeneous protein is vitreously frozen in a thin film of solution to form a glass-like ice, which is then imaged under cryogenic temperatures (−170 °C) in a transmission electron microscope (TEM) [6]. Many cryoEM micrographs are collected, with protein “particles” imaged in different orientations in 2D. Many 2D particle views (also known as “projections”) are collected and used to reconstruct an atomic 3D model [7] by iterative alignment, classification and averaging. The resulting micrographs are noisy; the low signal is due to a number of causes - microscope-related aberrations, low doses of electron exposure applied to the radiation sensitive protein specimen [8], movement of the specimen upon imaging [9], and due to the process of high resolution image formation in the TEM [10] [11]. Thus, the protein particles of interest are sometimes challenging to identify in these 2D cryoEM micrographs, especially certain orientations of the particles. Several software solutions have been developed to address this problem. These solutions require a large number of particles to estimate accurately the relative angular orientations of these protein particles in 3D, which is used to create 3D reconstructions of averaged protein particle structures.

One solution, RELION [12], uses the Bayesian approach and expectation maximization algorithm to solve for protein structures. RELION’s workflow allows users to manually identify and select (“picking”) particles in 2D cryoEM micrographs; then aligns, classifies, and averages those particles to make 2D class averages. Suitable 2D class averages are manually selected as templates, which are then used to perform automated particle selection (“autopicking”) by assessing image blocks’ correlation to the template particles [2]. EMAN2 [13, 14] is another popular solution to create 3D reconstructions from cryoEM micrographs. EMAN offers users three methods to pick particles from manual samples: local search, region search, and a simple deep neural network for pattern recognition based on THEANO [15]. One recent software called cryoSPARC [16] has a very promising approach for picking particles whereby the algorithm searches for a global optimal model from the regularized likelihood function instead of using an expectation maximization algorithm.

Although these software solutions have different methods for the particle picking process, the common point is that template particles (a set of good particles selected by experts from noisy cryo-EM micrographs also known as the "ground truth") needs to exist. This manual picking task is time consuming, and often an expert is not available. After obtaining good 2D class average templates, these software packages rely on two main methods to automatically select particles: template matching by cross-correlation or pattern recognition by a simple deep neural network. Template matching is sensitive to noise and may obtain strong bias [2], whereas neural networks are normally less sensitive and usually perform better. Unfortunately, simple neural networks are not robust enough to handle all the complicated noise patterns or the illumination/transmission variations in cryoEM micrographs. Also, necessary particle orientations cannot be usually obtained with the stringent constraints required (such as high picking threshold) due to the low contrast nature of cryoEM data. If we implement low picking thresholds, it is likely that these programs tend to pick not only particles, but also noise patterns known as false positives [17]. Unfortunately, one has to compromise to have either picking accuracy or false positive avoidance. Development of machine learning, specifically deep learning, holds much promise to improve and automate biomedical image analysis [18, 19, 20].

For microscopic image analysis, [21] many machine learning approaches have been utilized from support vector machines (SVM) to convolutional neural networks (CNN). A recent study of Shin *et al*. [22] has employed deep learning models to learn semantics in MRI scans, and to extract features to detect different organs. Xie [23] proposed a novel deep neural network for robust nucleus localization, where, instead of using a pixel-wise classier or a regressor, they combined CNN with a nonlinear voting transformation. Inspired by the success of these examples, we sought to apply these deep learning methods to cryoEM image analysis, specifically particle picking, which is the most tedious step in most cryoEM image analysis workflows.

In this paper, we introduce a Deep Regression Picker Network (DRPnet) to predict a probability map of particle centers and then classify those candidates to pick particles. The paper is organized as follows: in Section 2, we review recent advanced methods used to detect particles in microscopy images. Then, we explain the methods we used in this study and how DRPnet works in Section 3. In the final section, we present our experimental particle detection results for EMPIAR-deposited datasets, and 3D reconstructions obtained using these particles. Our major finding is that DRPnet finds particles with greater angular orientation coverage, and the certainty of the angular assignment is improved, as compared to those generated with RELION[12] template based autopicking [2]. This is especially helpful if the dataset is plagued by low particle numbers or the particle symmetry is low or unknown. Single particle analysis 3D reconstruction results using particles picked with DRPnet show improved resolution as compared to results obtained from 3D reconstruction using RELION software [24], without required network training or a dependence on particle size.

## 2 Related Work

DeepPicker [25] is a fully automated particle picking program based on deep neural networks. Before DeepPicker, other packages such as RELION and EMAN2 used semi-automated algorithms and workflows to collect particles. DeepPicker slides a window, with the same size as a specified particle (“box”), across a micrograph with a default step size of four pixels to collect candidate patches. All image patches are normalized and fed into a convolutional neural network to determine whether they belong to a qualified class of particles or not. In the fully automated mode, DeepPicker has a pretrained network to pick particles for the first time as positive training samples. The negative samples are randomly selected far from the positive samples using a spatial distance threshold. All of those samples are then used to train a fresh convolutional network to pick the final particles. However, in our experiment with *β*-galactosidase dataset (EMPIAR-10017), the pretrained mode did not obtain a satisfactory result. Next, we worked through DeepPicker’s training mode. In training mode, DeepPicker can adopt particles from RELION (using a manually picked .star file) to train the DeepPicker network. This second approach provided suitable results with higher particle picking accuracy and better capture of different particle orientations. In summary, DeepPicker worked quite well in the semi-automated mode. However, in fully automated mode, because of its simple four layer network configuration, DeepPicker was not powerful enough to acquire appropriate results with low contrast particles in EMPIAR-10061.

Zhu *et al*. [17] introduced a convolutional neural network (CNN) composed of eight layers called DeepEM to pick particles. DeepEM has three convolutional layers, one fully connected layer, and it requires data augmentation for both learning process and recognition process. The experiment was performed on a keyhole limpet haemocyanin (KLH) dataset and showed improved detection performance avoiding unwanted particles and non-particles. This framework, however, was only tested on one dataset (KLH; collected with a Tecnai F30 Twin on a Gatan Ultrascan 4000 CCD camera). If one required a fully automated detection framework, this simple network needs to be more versatile to adapt to a variety of differently shaped particles.

Xiao and Yang [26] employed Fast-RCNN [27] to select particles from micrographs. Instead of using a default region proposal function for Fast-RCNN, they made use of a sliding window to provide candidate regions for the detection CNN. This approach is equivalent to regular CNN architectures as the one used in DeepPicker. The strength of this method is that the network was trained with three classes instead of two, in order to discriminate false positive samples.

Heimowitza *et al*. [28] introduced a new picker called APPLE picker based on cross-correlation. It automatically selects reference windows to compare and eliminate the need to prepare manual templates. Compared to RELION’s manual template generation, APPLE picker works faster, accurately, and automatically. However, APPLE picker did not work well with high concentrations and hollow cylindrical particles (KLH, GroEL).

crYOLO [29] utilized the first generation of YOLO network [30] to detect particles. YOLO is based on a fixed-grid regression method. Compared to other region-based methods, this fixed-grid helps YOLO to work faster in real-time [31]. However, YOLO have detection problems with small objects and objects with unusual aspect ratios, and also suffers from high localization errors [31]. crYOLO performed detection very fast, up to six micrographs per second with image dimensions of 1024 × 1024 pixels [29]. crYOLO attained higher accuracy (recall, precision, and AUC) compared to the original YOLO model when working with small objects as particles in those test datasets. Despite modifications to the original YOLO architecture, the experiments do not mention how the crYOLO architecture performs using particles of other sizes and aspect ratios.

Most recently, two more packages, WARP and TOPAZ, are gaining popularity. *WARP* [32] uses a picking function based on BoxNet [33] for cryo-EM data processing. This deep neural network was trained with multiple EMPIAR raw and simulated data. As a result, it can pick many types of particles and can mask out the artifact regions. WARP supports network re-training with custom data and runs on a Window platform for integration with the cryoEM pipeline of data collection, preprocessing, particle picking and single particle analysis. TOPAZ [34] is another particle picking program using deep learning. Similar to DeepPicker [25], it examines micrograph patches and uses non-maxima suppression to select only the patches with highest scores as the most likely particle instances. One major difference between these approaches is that TOPAZ was trained with positive and unlabeled samples, instead of positive and negative samples.

Since a robust and flexible deep learning tool had not been developed to date for cryoEM particle picking, we set forth to develop a deep learning-based particle picking system that would work on multiple types of data (vitreous ice and negative stain), with various particle sizes and aspect ratios, and collected using different detectors (direct detection and CCD) with different defocus ranges. Below, we present our work on “Deep Regression” particle picking (Deep Regression Picker Network; DRPnet, Figure 1). The proposed system works on multiple types of data (tested on four protein datasets), from various detectors, with improved speeds (testing time is approximately ~6 sec/micrograph on Nvidia GTX 1080 GPU with 8GB memory and SSD storage) compared to manual template selection (~30 min to select 1000 particles) required by RELION. The outputs of DRPnet were compared with the outputs of the RELION autopicking and produced final maps with similar resolution.

**Figure 1:**
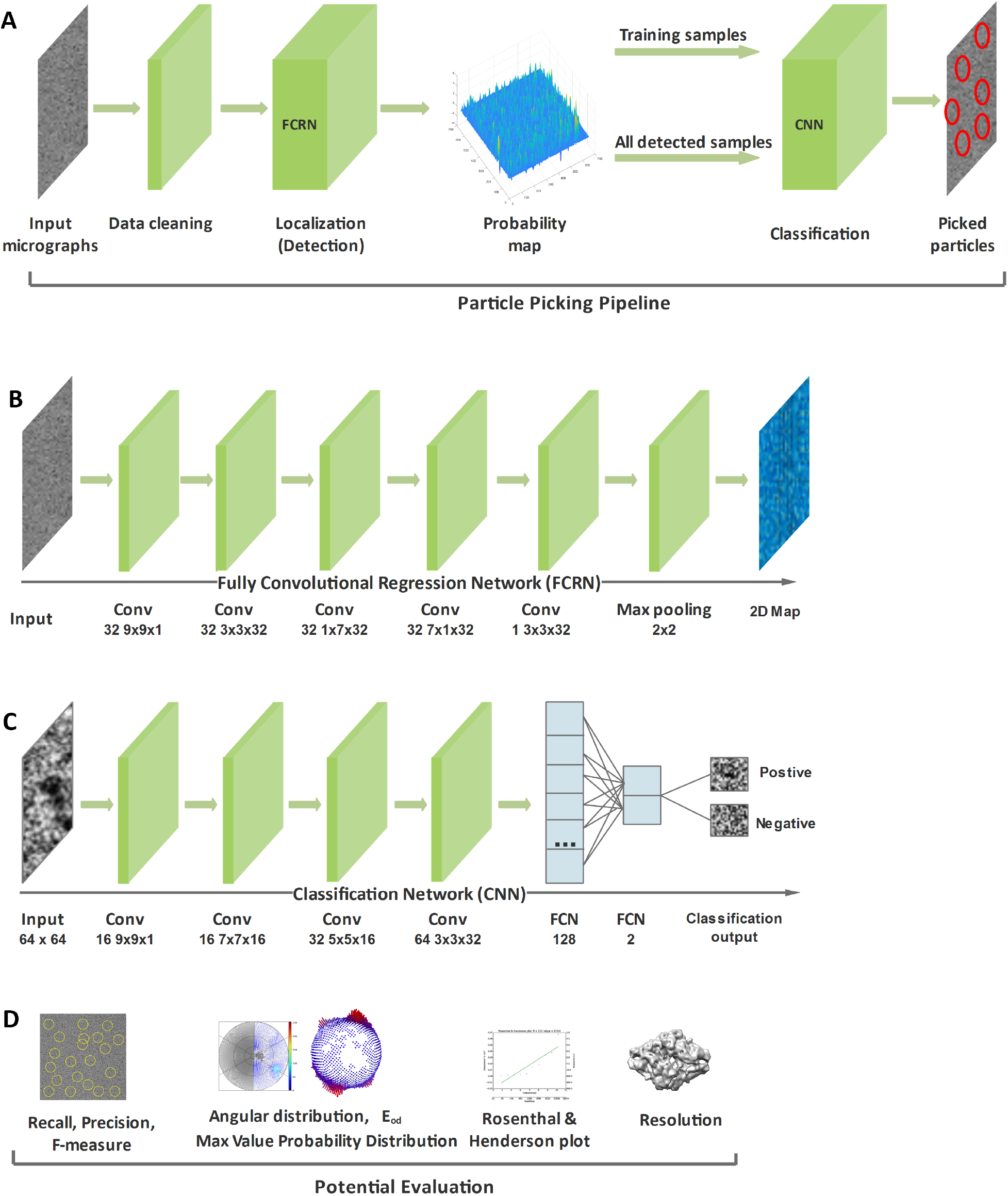
The cascade architecture of the Deep Regression Picker Network (DRPnet). (A) Overview of particle picking pipeline of DRPnet, from input micrographs to picked particles. (B) Architecture of Fully Convolutional Regression Network (FCRN) for initial particle detection. (C) Architecture of Convolutional Neural Network (CNN) for refining classification. (D) Potential useful evaluation metrics.

## 3 Material and Methods

### 3.1 Development of DRPNet Particle Picking Pipeline

Particle picking process in DRPnet involves two steps: localization (detection) and classification. In classical computer vision, object detection and classification processes often rely on carefully hand-crafted image features and descriptors such as HOG (Histogram of Oriented Gradients) [35], SIFT (Scale-Invariant Feature Transform) [36], or SURF (Speeded-Up Robust Features) [37] etc. that are extracted from image patches. These descriptors are then fed to supervised machine learning models such as SVM (support vector machine), random forests, and neural networks to cluster or to classify those feature vectors. Given an input image, a scanning window slides over the image and independently classifies each image patch based on features extracted from the patch. The success of these systems heavily depends on developer selected features that may not capture complexities of the underlying visual patterns or adapt to new patterns.

On the other side, deep learning approaches rely on data itself to identify discriminative features to perform the given task. Convolutional neural networks (CNN), a class of deep learning methods heavily used for image analysis, extract features over multiple convolutional layers with different sizes and different numbers of filters whose coefficients are learned during training. Earlier layers of convolutional networks learn deep features, while later fully connected layers perform the classification or regression tasks. Deep detection approaches can be coarsely grouped as single-stage and two-stage. Single-stage detectors such as YOLO [38] and SSD [39] rely on regular dense sampling of objects, scales, and aspect ratios and perform detection and classification in one step. Two-stage detectors such as FasterRCNN [40] and Mask-RCNN [41] first produce a sparse set of candidates (region proposals). These candidates are then classified into a number of classes, foreground versus background in our case. Faster-RCNN and Mask-RCNN have shown great performance in detection and classification tasks on image datasets such as Pascal VOC [42] and MS COCO [43]. However, these models were originally designed to work with color (RGB, HSV, CYMK) images and larger objects with rich color, texture, and shape features. Unlike color images and objects in these datasets, electron micrographs are grayscale, low contrast, and may contain noise and artifacts (support-film edges and surface ice contamination). We required more versatile network than Faster-RCNN’s Region Proposal Network (RPN) [40] to perform automated particle detection in cryoEM micrographs. In order to overcome the aforementioned issues, we propose the Deep Regression Picker Network (“DRPnet”, Figure 1), which is based on the blob detection concept [44]. After the initial training with one dataset, the proposed DRPnet can work with multiple particles having various sizes and shapes from different datasets without need for retraining. We also realized that using scanning window and a single network for both localization and classification limits the ability to recognize patterns and/or discriminate different types of false positive patterns. For this reason, our particle picking pipeline (Figure 1A) includes two steps: localization (detection) relying on a fully convolutional regression network (FCRN) (Figure 1B) and classification relying on a classification convolutional neural network (Figure 1C). Although we used a supervised CNN model to classify true versus false positives, training data for this classification network were collected in an unsupervised manner. To identify true versus false positives in a semi-supervised manner, we used FCRN’s measure of confidence where high level corresponded to a true positive and low level corresponded to a true negative. The initial network was trained using the EMPIAR-10005 dataset [1]. This scheme allows DRPnet to operate on other datasets automatically from detection through classification.

Our particle picking pipeline also included a micrograph preprocessing step to enhance contrast and to correct transmission/illumination artifacts in order to ensure optimal performance by both networks. Picked particles were then fed to RELION [45] software for 3D reconstruction. Processing steps involved in the proposed automatic particle picking pipeline are described in the following subsections.

### 3.2 Data Preprocessing

#### 3.2.1 Data Cleaning: Illumination/Transmission Correction

Illumination (beam centering) and/or electron transmission (transmitted electron signal) can misguide DRPnet’s detection because particle picking significantly relies on intensity of grayscale images. To obtain the best DRPnet detection performance, it was necessary to reduce the influence of illumination/transmission variations. We compute local averages of illumination by applying a low-pass filter with a very large kernel. The corrected intensity value of each pixel is obtained by subtracting local and adding global intensity averages, as in Equation 1:

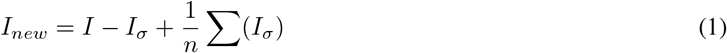

where *I* is the original image, *n* is the number of pixels in the image, and *I_σ_* is image smoothed with a Gaussian filter of sigma *σ*, set heuristically according to particle size. Very low contrast and extremely high noise characteristics, make electron micrographs more challenging compared to typical color or grayscale images.

#### 3.2.2 Distance Transform

Distance transform of a binary image is a grayscale map where each pixel’s value represents the distance to the closest boundary. It is calculated at each pixel by the distance to the nearest background pixel. In this study, we applied distance transform to the binary mask of particles to produce a representation of a probability map of the particle centers (Figure 2B).

**Figure 2:**
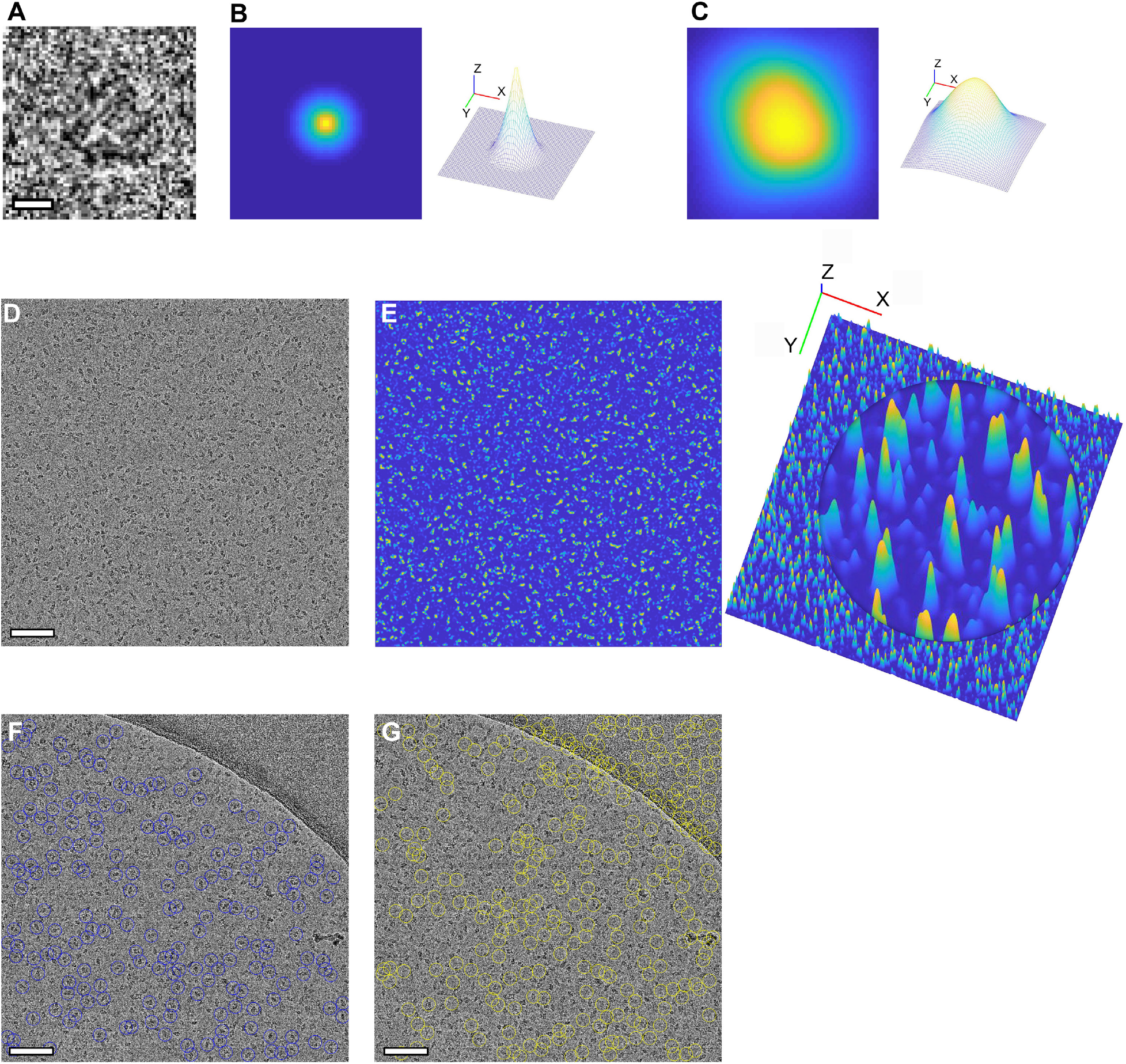
Training and test examples of DRPnet on cryoEM micrographs. (A) Sample input particle from cryoEM micrograph of TPRV1 (scale is 50 Å). (B) Corresponding ground truth training label (left) and its 3D visualization (right), obtained by applying distance transform to the binary particle mask - with blue and yellow indicating lower and higher distance values respectively. (C) Smoothed, 2D predicted map (output of the fully convolutional regression network shown in Figure 1B) for a single particle (left) and its 3D visualization (right). (D) Example of original input cryoEM micrograph (scale is 500 Å). (E) Predicted (2D) map of entire cryoEM micrograph by DRPnet (left), and 3D predicted map visualization with circled area showing a magnified view of the maxima (yellow-capped peaks) in panel E. (F-G) Positive (F, blue circles) and negative (G, yellow circles) particles used to train the classification network shown in Figure 1C. Positive samples are true targets, and negative samples represent false targets. These positive and negative training samples are selected in an unsupervised way using the prediction confidence values from the fully convolutional regression network depicted in Figure 1B with high and low confidence particles corresponding to positive and negative training samples respectively (scale is 500 Å).

### 3.3 Deep learning

#### 3.3.1 Fully Convolutional Regression Network (FCRN)

To address the challenges of automated particle picking, we have treated this as a blob detection problem - where each particle is roughly a convex blob with texture different than the background (Figure 1B & 2A). Additionally, we assume that the dimensions and shapes of particles are relatively similar. Given a rough size estimate, our goal is to locate particle centers. For the purposes of this study, segmentation of particle regions from image background is unnecessary. Our proposed model is a Fully Convolutional Regression Network (FCRN), trained to predict particle centers by producing a likelihood map where local maxima correspond to the particle centers (Figure 2). Given the ground truth particle centers (as provided by the datasets in EMPIAR) along with a particle size estimate, first, binary particle versus background masks were produced for the cryoelectron micrographs. Then, distance transforms of those binary masks are then used as training labels for the proposed FCRN model (Figure 2A). The distance transform for the background pixels are set to zero (Figure 2B). whereas the values of the distance transforms for the foreground regions (or the particles) increase from the particle’s boundaries towards particle’s centers, reaching their local maxima at the particle centers (Figure 2B). Therefore, a continuous regression model (FCRN), rather than discrete classification models, is used to localize particle centers. Because of its continuous nature, this model enables localization of individual particles within dense clusters, and ensures better scale adaptation, which allows detection of different sized particles without need for retraining the network. This important feature is missing in the existing algorithms reported to date. Furthermore, since the proposed network is built without fully connected layers, any size micrographs can be used as input to the network. The proposed FCRN has seven layers (Figure 1B), one input, five convolutional, and one max-pooling layers [46]. With its simple structure, our approach has smaller computational cost and training data needs, compared to more complex deep learning models such as Fast-RCNN.

The proposed network was implemented using Matlab deep learning toolbox. The network was trained with image patches from raw micrographs and associated training labels, centered and cropped on particle regions. These image patches are of the same size as the FCRN input layer (Figure 2B). The number of patches extracted from each image is equal to the number of particles in those images. The network is trained using the Adam optimizer [47]. Since our network does not have fully connected layers, it does not require fixed sizes of input images. In the testing stage, we directly applied the trained FCRN model to different-sized input images (Figure 2D) to predict associated particle center probability maps (Figure 2E).

#### 3.3.2 Classification Network

As shown in the working pipeline in Figure 1B, the FCRN probability map output indicates locations of all detected particles. The second convolutional neural network (CNN) classifies these particles for the final keep-or-delete selection process 1C. We extracted two subsets from the particles detected by the first FCRN network, and trained a second two-class classification network using stochastic gradient descent optimization [46]. Positive training samples were particles detected with high confidence, further refined by removing those with low standard deviation of intensity. Negative training samples were particles with low probability in the detection map as well as particles which had a low standard deviation of intensity. Negative particles included patches of background, carbon edges, ice contaminated regions, and weak patterns. Using the second convolutional network, these particles are classified into positive particles (Figure 2F) and negative particles (Figure 2G). This second classification network learns texture patterns of true particles versus false positives caused by various artifacts, and provides a refinement to the particle detection process in an efficient unsupervised manner. This network has four convolutional layers, with each followed by a max-pooling layer [46] for feature extraction and two fully connected layers [46] to decide whether to pick or discard a particle patch using the learned features.

### 3.4 Performance Metrics

In this research, we applied multiple metrics to measure our DRPnet-based particle picking performance (Figure 1D). The first metric uses recall, precision, and F-measure [48] to evaluate how accurately our model predicts, which can only be calculated if a ground truth exists. After picking particles, we use them to reconstruct 3D maps of protein structures to measure resolution, which serves as our second metric. Finally, we analyze angular distributions of those 3D reconstructions and calculate the efficiency E_od_ [49] to understand the effect of orientations of the final 3D reconstruction resolution from those particles picked by either DRPnet or RELION. We also visualize the effect of number of particles on structures’ resolution by Rosenthal-Henderson Plots [50] [51].

#### 3.4.1 Recall, Precision & F-measure

The overall detection performance of all particles are evaluated using three metrics: recall, precision, and F-measure [48]. Initially, we collected the detection results in a confusion matrix which has columns to indicate the ground truth (a real particle), and the rows to indicate prediction results. True Positive (TP) and True Negative (TN) are two cases where the prediction and ground truth match (Table 1). Otherwise, when prediction and ground truth are different, we will get a false positive (FP) if we detect more than the ground truth, or we will have false negative (FN) if we miss the ground truth cases. From these basis terms, Recall is defined as the number of true positives divided by the number of true positives plus the number of false negatives (Equation 2). Recall indicates the ability to catch all the ground truth cases. Precision is defined as the number of true positives divided by the number of true positives plus the number of false positives. Precision indicates how the prediction is relevant to the ground truth (Equation 3). In order to detect all the ground truth particles, one may decide to return all the images patches. In which case, recall would be high but precision would be low because only a small portion of the returned image patches would correspond to true particles. Conversely, if one only selects the high confidence predictions, precision would be high, but would result in low recall scores due to some missing ground truth cases. F-measure, computed as two times of product of Recall and Precision, then divided by sum of Recall and Precision, is useful to assess our model’s prediction (Equation 4).

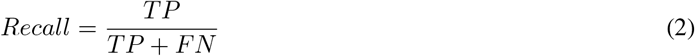

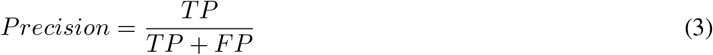

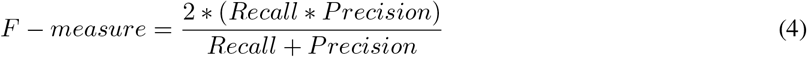

**Table 1:** Prediction Outcomes

|  | Actual Positive | Actual Negative |
| --- | --- | --- |
| Predict Positive | TP | FP |
| Predict Negative | TN | FN |

#### 3.4.2 Resolution

The second evaluation metric we applied to evaluate the performance of particle picking is the final resolution of the 3D reconstruction. In cryo-electron microscopy, we rely on the concept of signal to noise ratio, and measure the internal consistency of of the 3D reconstructions generated by each particle picking method (DRPNet or Relion’s template based method). We take advantage of some characteristics of Fourier transform: correlation between 3D reconstructions can be represented by a product in Fourier space, and then we split that product into shells by radial frequency to compute Fourier Shell Correlation (FSC; Equation 5) [52, 53, 12, 54].

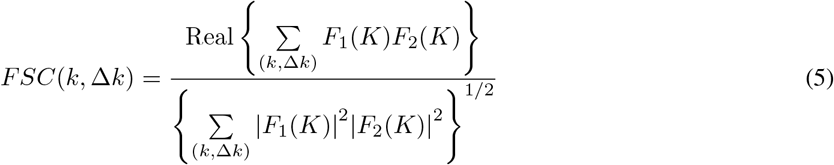

*K*: spatial frequency vector

*k* = |*K*|, is the size of spatial frequency

Δ*K*: ring width or shell thickness

*F*_1_(*K*), *F*_2_(*K*): Fourier transforms of two half set reconstruction

A plot of FSC vs. spatial frequency depicts their relationship as we showed in Figure 6C, 6F and 6I. Besides that, the connection between FSC and Spectral Signal to Noise Ratio (SSNR) is explained by [55] as following:

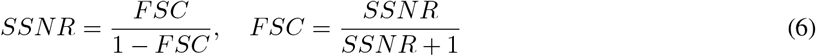

FSC is a biased estimate of SSNR. For a large number of images: Variance(SSNR) = Variance(FSC), and the bias is negligible. When FSC is calculated for a data set which is split into halves, the relationship is:

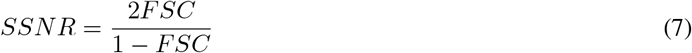

With the relationship between FSC and SSNR, we can use FSC to measure the internal consistency of our 3D reconstruction. We can correlate a a specific level of FSC to an equivalent spatial frequency. That frequency has a unit of 1/Angstroms (1/Å) which is the inverse of the second measured metric, resolution. In order to divide our data into two volumes to calculate FSC, we can select odd particles for the first 3D reconstruction and the even particles for the second reconstruction after we construct the 3D reconstruction with all particles. This way is very simple, but the two halves of our reconstructions are not independent. Another way is splitting particles from the beginning and build two 3D models. This method is often referred as “Gold standard Fourier Shell Correlation” [56]. A cut-off level is necessary to identify structures’ resolution. A reasonable criterion includes only Fourier information that is above the noise level. Typical cut-off level for Gold standard split, as used in Relion, is 0.143 [57, 12, 56] (Figure 6C, 6F and 6I), and for odd-even split is 0.5 [58, 12]

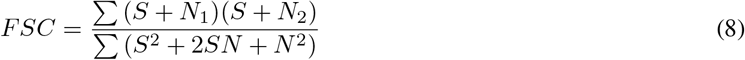

where *S* denotes the signal and *N* = *N*_1_ = *N*_2_ are noise in half datasets.

#### 3.4.3 Rosenthal-Henderson Plots

As originally proposed by Rosenthal-Henderson [50], plotting of the inverse-squared resolution as a function of the number of particles allows us to compare numbers of particles picked with DRPnet or RELION and the effect of particle numbers on 3D reconstruction resolution [51].

#### 3.4.4 Angular distribution

Besides two metrics above, we analyzed the angular distributions recorded in RELION .star files of 3D Refinement jobs to understand the difference in orientation of particles picked by DRPnet and RELION’s Autopick. Those distributions showed the effect of orientation on resolution and clarified the difference of final resolution of structures by DRPnet and RELION. We visualized the angular distribution by 2D scatter plots of rot-tilt angles [59], and visualized the bild files of 3D refinement jobs by 3D histograms in Chimera [60]. Furthermore, we employed efficiency number E_od_ [49] to confirm the angular distributions’ effect. By using corresponding point spread function, Naydenova and Russo [49] computed E_od_ to assess how the angular distribution contributes to the reconstruction results. An E_od_ value of 0.8 1 indicates a good orientation distribution and a uniform Fourier space coverage. Otherwise, if E_od_ is lower than 0.5, the orientation distribution is not effective to 3D reconstruction. Finally we plotted the “Max Value Probability Distribution”, an output from RELION’s 3D Refinement, where values close to one indicate that the probability distribution over all orientations and classes have converged to near-delta functions. We use this metric to provide confidence that RELION was accurately assigning orientations to the particles.

### 3.5 Overview of Methods

Taking cryoEM datasets available in EMPIAR [61]or locally generated negative stain micrographs, we utilized both DRPnet-particle picking or RELION (v3.0) autopicking to generate datasets with similar numbers of particles. For each EMPIAR data utilized, RELION requires generation of a manually picked particle set, usually around 1000 particles total. Particles used in our RELION comparison results were generated by randomly selecting a subset (1000) of particles deposited in EMPIAR that contributed to the final, refined high resolution 3D reconstructions previously published. These randomly selected particles can be considered the “manually picked” particles. These randomly-selected, manually picked particles were then processed through the standard RELION pipeline (template generation by 2D classification). DRPnet particles were generated as described in Sections 3.2–3.4. Both DRPnet particles and the RELION’s autopicked particles were then processed through RELION’s 2D classification to identify good 2D class averages (see Figure 2 in chapter 6 of Methods in Enzymology [62]). Both DRPnet and RELION-autopicked 2D class average templates were further 3D classified (allowing 5 possible classes) using the corresponding deposited, low-resolution (60 Å) filtered 3D reconstruction as the initial model. All good 3D classes were selected and refined to generate the final 3D reconstruction. RELION used gold standard as cut off level to calculate those structures’ resolutions.

## 4 Results and Discussion

### 4.1 Datasets

To demonstrate our model’s ability of working with multiple types of particles, we prepared a pretrained DRPnet model from one dataset and tested it on other datasets. We downloaded three cryoEM datasets from EMPIAR [63] including TRPV1 (EMPIAR-10005) [1], *β*-galactosidase (EMPIAR-10017) [2], and *β*-galactosidase (EMPIAR-10061) [3]. CryoEM datasets were acquired using an 300 kV accelerating voltage on FEI Titan Krios cryo-transmission electron microscopes or on a FEI Tecnai F30 Polara. For training DRPNet, we selected TRPV1 [1]. For a summary of these datasets, please see Table 2.

**Table 2:** Experimental datasets from EMPIAR used for DRPnet testing

| Dataset | Voltage (kV) | Camera | Type | Defocus Range ($\mu\text{m}$) | # Total Images | # Test Images | # Picked Particles |
| --- | --- | --- | --- | --- | --- | --- | --- |
| EMPIAR-10005 (TRPV1) | 300 | Gatan K2 | Vitreous | 1.5 - 3.5 | 871 | 661 (subset) | 60,000 |
| EMPIAR-10017 (β-gal) | 300 | FEI Falcon 2 | Vitreous | 1.0 - 3.0 | 84 | 84 (full set) | 50,000 |
| EMPIAR-10061 (β-gal) | 300 | Gatan K2 Summit GIF K2 | Vitreous | 0.6 - 3.0 | 1538 | 661 (subset) | 60,000 |

### 4.2 Training on EMPIAR-10005 (TRPV1)

To train the first FCRN network of DRPnet (Figure 1B) for particle detection, we used a subset of TRPV1 data (EMPIAR-10005), including 50 TRPV1 micrographs and 9751 ground truth particles extracted from deposited coordinates used for the high resolution 3D reconstruction [1]. These cryoEM micrographs have dimensions of 3710×3710 pixels and a pixel size of 1.2156 Å. We scaled them down by 3x to fit the scale of EMPIAR-deposited particle coordinates, and then used a box size of 64×64 pixels to extract particle patches. Those deposited particle coordinates were used to generate binary masks, then the distance transforms of those masks were calculated (Figure 2B) and used as the ground truth to train the FCRN network. The second classification CNN of DRPnet (Figure 1C) was trained to detect positive or negative training samples (Figure 2G-H), after testing the first FCRN network.

### 4.3 Testing on EMPIAR-deposited datasets

#### 4.3.1 Test on EMPIAR-10005 (TRPV1)

In order to validate our pretrained DRPnet model with TRPV1, we tested it on a set of 661 TRPV1 micrographs, collected on a Gatan K2 direct detector. TRPV1 micrographs have low contrast, and some do not contain any particles. We set the first network (FCRN) at normal detection level (sigma = 9, threshold = 2*5) to detect particles. To discriminate true positive and false positive detections, we select classification training samples at positive level (sigma = 9, threshold = 3*5) and negative level (sigma = 9, threshold = 1*5) to train the second network. As a result, DRPnet picked 61,282 particles. For RELION’s autopick, we selected sigma = 1 and threshold = 0.475 to pick 61,599 particles. We extracted particles with box size 200×200 pixels without scaling to perform the subsequent jobs. Taking both sets of picked particles forward through the RELION 3.0 pipeline, we selected 23,147 good particles picked by DRPnet and 22,830 good particles picked by RELION autopick.

Inspecting the same cryoEM micrograph picked with DRPNet (Figure 3A) or Relion’s template based autopicking (Figure 3B), the particles that are close to each other are able to be better picked using DRPNet (as discussed in Methods). Also, when inspecting the classes for DRPNet generated particles (Figure 3C) compared to Relion (Figure 3D) there are more “good” classes obtained with DRPNet (49) versus RELION (32). In this experiment, the resolution of DRPnet structure was 3.92 Å, equal to the resolution of RELION autopick structure which was also 3.92 Å(Figure 6C). Increasing the number of EMPIAR-10005 particles from 600 to 20,000, a Rosenthal & Henderson plot (Figure 6A) reveals that DRPnet produces a better 3D reconstruction resolution than RELION’s autopick, especially if symmetry is not imposed (C1).

**Figure 3:**
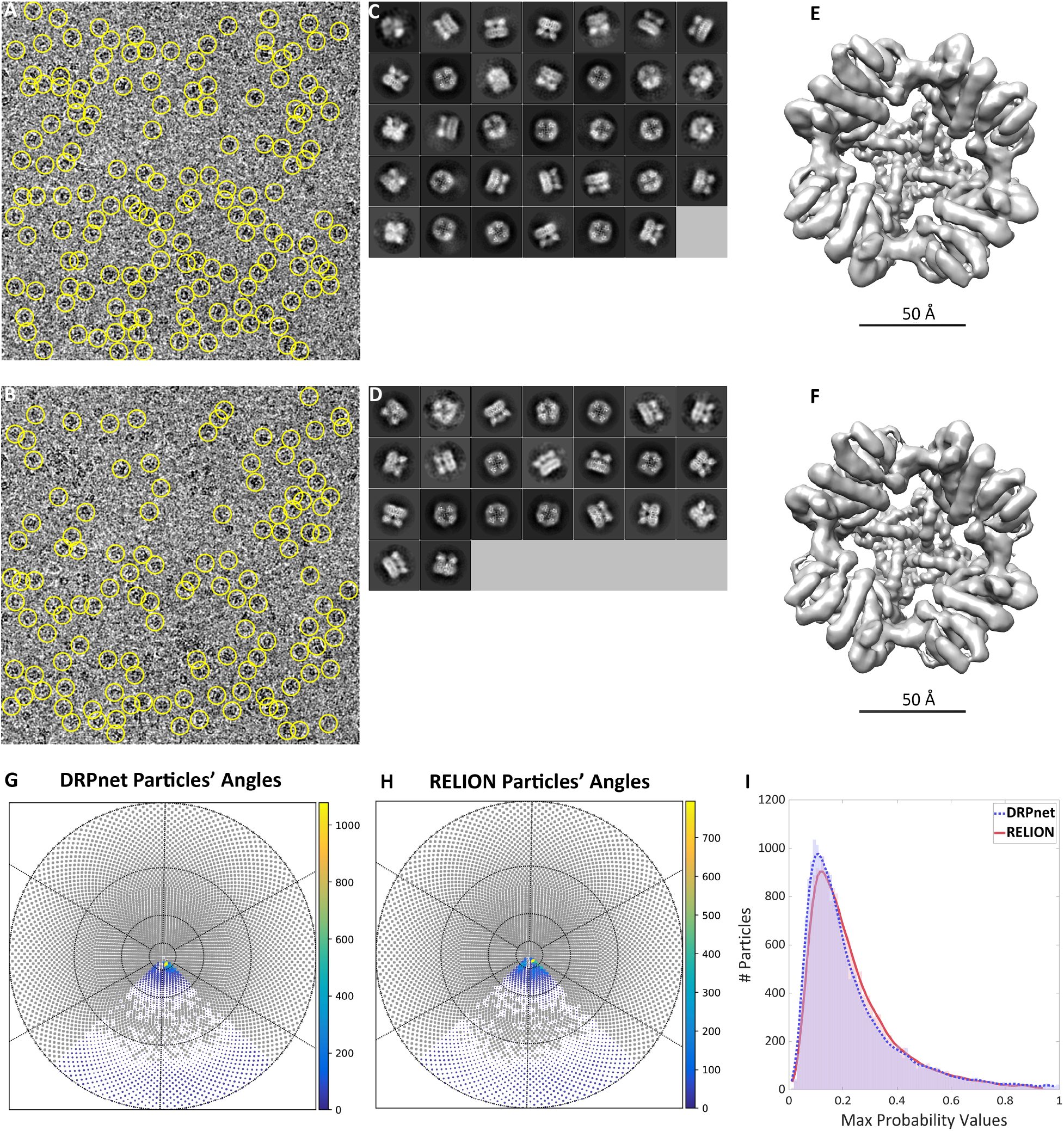
EMPIAR-10005 (TRPV1) testing of DRPNet. Representative cryoEM micrograph shows picked particle output (yellow circles) by DRPnet (A) and RELION (B). All picked particles were classified using RELION to select quality 2D class averages as displayed for DRPnet (C) and RELION (D). Then those particles were used to generate 3D reconstructions, DRPnet’s result (E) and RELION’s result (F), are superimposed, and displayed in gray isosurface respectively. The particles’ angular distributions were determined and refined during 3D reconstruction and are displayed in Rot-Tilt angle scatter plots for DRPnet (G) and RELION (H). A gray filled circle indicates a missing view. Overlay of the histograms of Max Value Probability Distribution (I) of particles picked by either DRPnet (blue dashed line) or RELION (red solid line).

We also studied the angular distribution to further analyze 3D reconstruction resolution. The histograms of Rot-Tilt angles (Figure 3G and 3H) and Table 3 show that the distribution of particle orientations with RELION’s autopicked particles were lacking compared to those picked with DRPnet. This result also agreed with the visualization of RELION’s bild files (Figure 6B), and the efficiency [49] E_od_ reported Table 3, with DRPnet having a E_od_ of 0.57 while RELION had E_od_ of 0.51. We imposed symmetry C4 for the TRPV1 3D reconstructions, however without imposing symmetry (Table 3, indicated with parentheses), we note that DRPnet picked particles had improvements 3D orientation coverage, which was further confirmed with improved E_od_, upper lines in the Rosenthal Henderson plots (Figure 6A) and resolution (Figure 6C) than RELION autopick. Besides the orientation angles, we evaluated the Max Value Probability Distribution to see how confident RELION was of the picked particles’ orientations in the 3D reconstruction, with a low Max Probability Distribution value indicating low confidence of orientation. As shown in Figure 3I, probability values of DRPnet have a mean of 0.230, and a standard deviation of 0.173 while RELION has a mean of 0.237 and a standard deviation of 0.161. The histogram of Max Value Probability Distribution shows that DRPnet picked more particles with higher values (blue area under the curve) than RELION (red area under the curve), thus these particles could be more confidently assigned their angle in the final 3D reconstruction.

**Table 3:** Test results (values in parentheses were obtained without imposing symmetry)

| Dataset | Program | # Picked Particles | # Selected Particles | # Missing Views (in total 12,288 views) | $E_{od}$ | Resolution (Å) |
| --- | --- | --- | --- | --- | --- | --- |
| EMPIAR-10005 (TRPV1) | DRPnet | 61,282 | 23,147 | 10,164 (7,807) | 0.57 (0.47) | 3.92 (6.57) |
|  | RELION | 61,599 | 22,830 | 10,389 (8,591) | 0.51 (0.32) | 3.92 (6.75) |
| EMPIAR-10061 (β-gal) | DRPnet | 60,267 | 49,582 | 9,196 (4,019) | 0.82 (0.76) | 2.78 (3.02) |
|  | RELION | 61,597 | 49,570 | 9,288 (4,320) | 0.80 (0.74) | 2.81 (3.06) |
| EMPIAR-10017 (β-gal) | DRPnet | 49,604 | 41,914 | 9185 (2,250) | 0.69 (0.70) | 4.21 (4.72) |
|  | RELION | 49,855 | 40,669 | 9,250 (2,412) | 0.69 (0.70) | 4.26 (5.06) |

**Table 4:**
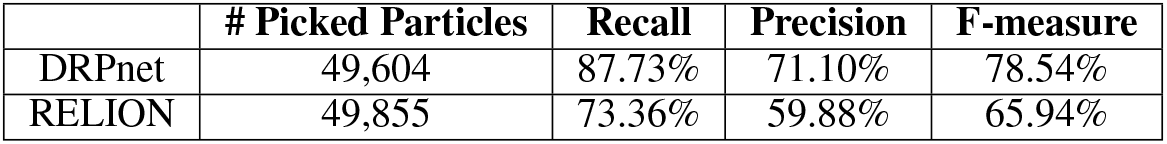
Recall, Precision and F-measure of EMPIAR-10017 (*β*-galactosidase)

|  | # Picked Particles | Recall | Precision | F-measure |
| --- | --- | --- | --- | --- |
| DRPnet | 49,604 | 87.73% | 71.10% | 78.54% |
| RELION | 49,855 | 73.36% | 59.88% | 65.94% |

Next, we tested on two other datasets to verify that DRPnet could not only select particles with diversified shapes, but also improved orientation ranges resulting in higher resolution structures.

#### 4.3.2 Test on EMPIAR-10061 (*β*-galactosidase)

In the second test, we picked particles from a selected 661 micrographs from EMPIAR-10061. This dataset was collected with little defocus and at high magnification, which resulted in extremely low particle contrast. These micrographs have a size of 7676×7420 pixels, and the original pixel size is 0.3187 Å. We set DRPnet to work at detection level (sigma = 1, threshold = 1*0.01) and scale factor of 1/8, and then trained classification network with positive samples as a subset of detected particles with high standard deviation (s > 0.25) and negative samples as a subset of detected particles with high standard deviation (s < 0.25). Accordingly, DRPnet picked 60,267 particles from those micrographs. For RELION’s autopick, we selected sigma =1.2 and threshold = 0.15 to pick 61,597 particles. All subsequent jobs including extraction, classification and refinement used a binning factor of 2, a box size of 384×384 pixels and a pixel size of 0.637 Å. In 3D structure refinement, we kept 49,592 good particles of DRPnet and 49,570 good particles of RELION autopick. The results showed that DRPnet structure had a resolution of 2.78 Å, while RELION structure had a resolution of 2.81 Å. Figure 4G and 4H illustrate the angular distribution of particles picked from dataset EMPIAR-10061 (*β*-galactosidase). The histograms of Rot-Tilt angles show that the particles of DRPnet (Figure 4G) missed less views than particles of RELION autopick. This result also agreed with the efficiency numbers E_od_ in Table 3 as well as the illustration of RELION’s bild files in Figure 6E. In our experiment, DRPnet had E_od_ of 0.82 while RELION had E_od_ of 0.80 with two-fold symmetry (D2). Without imposing symmetry (C1), the values in parentheses in Table 3 still show that DRPnet missed less views, and had better E_od_ and resolution than RELION autopick. Besides the angles, we evaluated the Max Value Probability Distribution to see how those picked particles fit best to the reconstruction process of RELION. The histogram of Max Value Probability Distribution (Figure 4I) shows that DRPnet had more higher values than RELION. Those probability values of DRPnet have a mean of 0.339, and a standard deviation of 0.201 while RELION has a mean of 0.324 and a standard deviation of 0.205. Rosenthal & Henderson plot in Figure 6B shows that when the number of EMPIAR-10061 particles increase from 1,200 to 38.000 particles, DRPnet obtained better resolution than RELION’s autopick in both cases with and without symmetry imposed.

**Figure 4:**
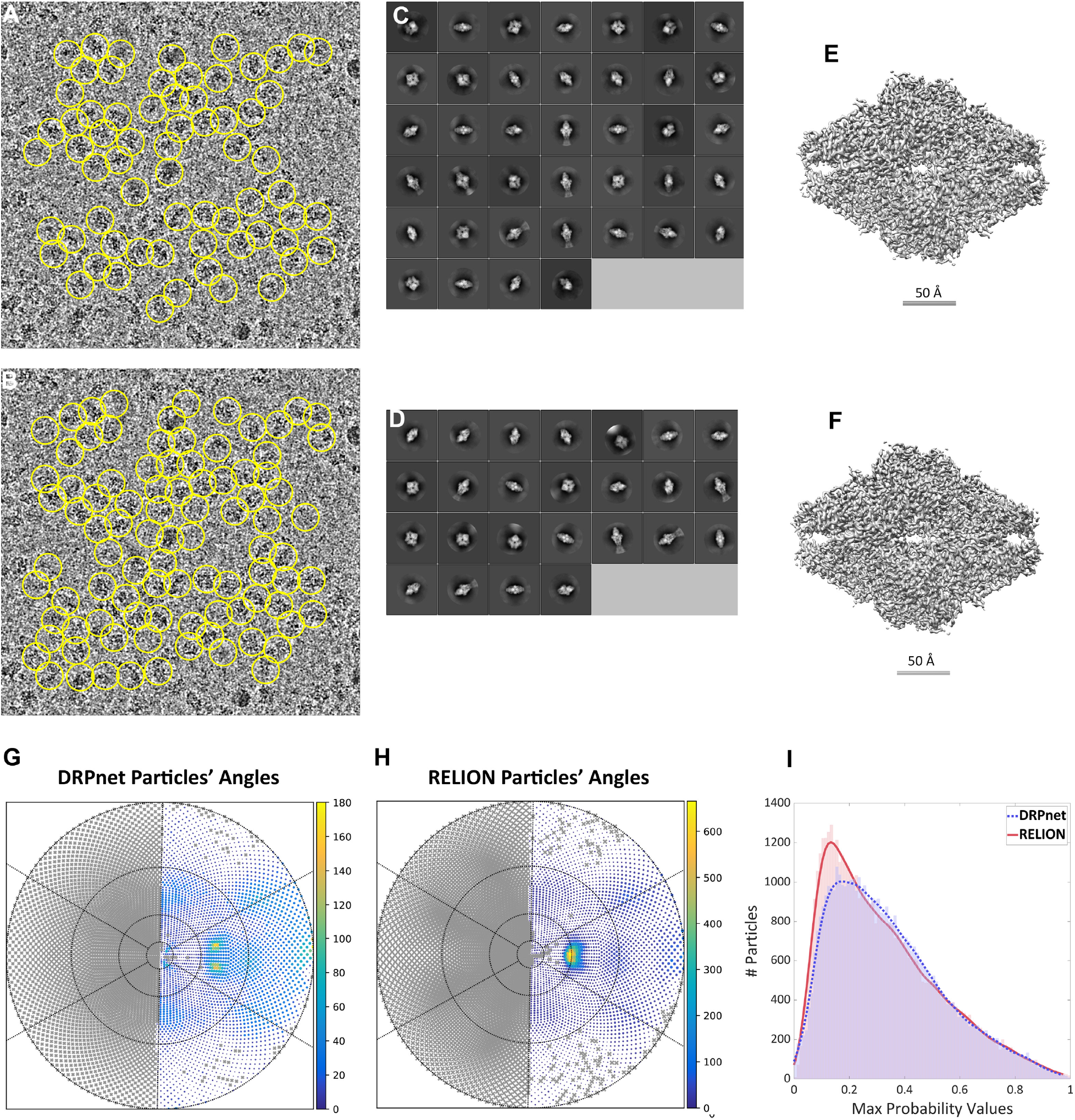
Test results of EMPIAR-10061 (*β*-galactosidase): The first two images are examples of picked particles by DRPnet (A) and RELION (B). All picked particles were classified using RELION to select quality 2D class averages as displayed for DRPnet (C) and RELION (D). Then those good particles were used for 3D reconstruction, and the isosurface superimposed renderings of the maps were generated from DRPnet’s result (E) and RELION’s result (F) respectively. The particles’ angular distributions in reconstruction process are displayed in Rot-Tilt angle scatter plots by results of DRPnet (G) and RELION (H). A gray “X” symbol indicates a missing view. The last sub-figure overlays the histogram of Max Value Probability Distribution (I) of particles picked by DRPnet (blue dashed line) and RELION (red solid line).

#### 4.3.3 Test on EMPIAR-10017 (*β*-galactosidase) data

After EMPIAR-10005 (TRPV1) and EMPIAR-10061 (*β*-galactosidase) test, we conducted the third test on entire EMPIAR-10017 (*β*-galactosidase) dataset. This dataset has an image size of 4096×4096 pixels and a pixel size of 1.77 Å. We set DRPnet to detect particles at picking level (sigma = 7, threshold = 2*7), and selected training samples for classification at positive level (sigma = 16, threshold = 3*7.5) and negative level (sigma = 12.5, threshold = 2*5) to pick 49,604 particles. On the other side, RELION’s autopick picked 49,855 particles. Those particles were extracted with a box size of 200×200 pixels at original scale for 3D map reconstruction later. With a provided ground truth (particles manually picked by an expert [2]), we could evaluate detection accuracy for this dataset. Table 3 shows that our DRPnet has recall of 87.73 %, and precision of 77.10 %, and F-measure of 78.54 %. Compared to RELION autopicking, DRPnet’s results are better when RELION used manually selected templates to pick particles. RELION achieved a recall of 73.36 %, precision of 59.88 %, and F-measure of 65.94 %. With the total number of picked particles around 50.000 particles from each algorithm, we run 2D classification to filter good particles. According to visual inspection, we kept only good average templates after 2D and 3D classification for 3D map reconstruction. The number of good particles corresponding to DRPnet’s pick and RELION’s autopick are 41,914 and 40,669, respectively. Our experiment showed that DRPnet structure had a resolution of 4.21 Å while RELION structure had a resolution of 4.26 Å. Figures 5G and 5H illustrate the angular distribution of particles picked from dataset EMPIAR-10017 (*β*-galactosidase). The histograms of Rot-Tilt angles show that the particles of DRPnet (Figure 4G) miss less views than particles of RELION autopick. In table 3, with two fold symmetry imposing (D2), DRPnet misses 9,185 views while RELION’s autopick misses 9,250 views. Without symmetry imposing (C1), the number of missing views between them is further increased to 162 views. Both DRPnet and RELION’s autopick gained the same number of E_od_ of 0.69 with two fold symmetry (D2), and 0.70 without symmetry (C1). Besides the view angles, the Max Value Probability Distribution shows us how those picked particles fit best to the 3D reconstruction process. The histogram of Max Value Probability Distribution (Figure 5I) indicates that DRPnet has more higher values than RELION. Those probability values of DRPnet have a mean of 0.27, and a standard deviation of 0.151 while RELION has a mean of 0.24 and a standard deviation of 0.136. Rosenthal & Henderson plot in Figure 6C shows that when the number of EMPIAR-10017 paritcles increase from 1,200 to 38,000 particles, DRPnet obtained better structure resolution than RELION’s autopick, especially in case of no symmetry (C1).

**Figure 5:**
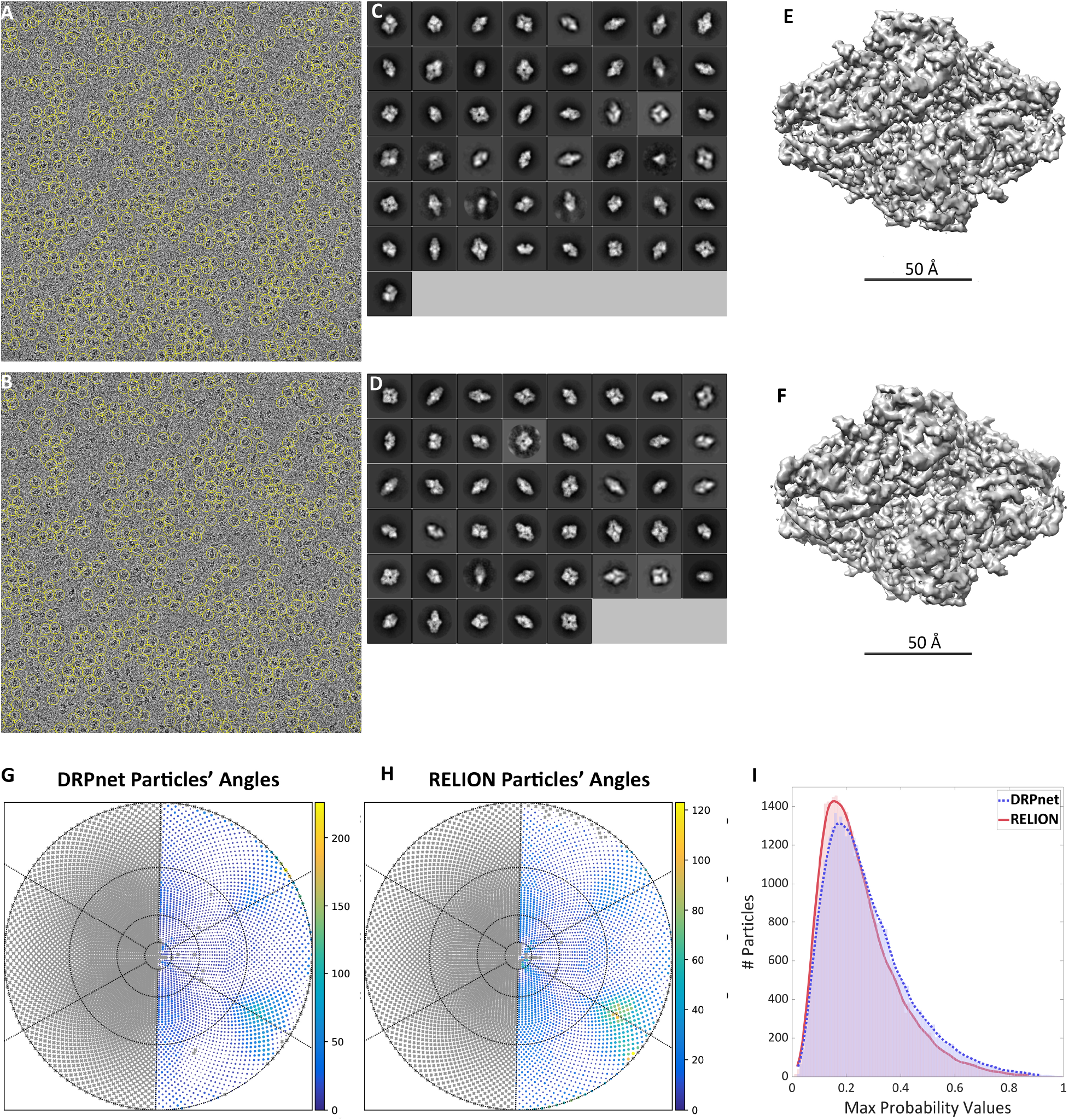
Test results of EMPIAR-10017 (*β*-galactosidase): Two examples of picked particles by DRPnet (A) and RELION (B). All picked particles were classified using RELION to select good 2D class averages as shown for DRPnet (C) and RELION (D). Then those good particles were used for 3D reconstruction, and the isosurface superimposed renderings of the maps were generated from DRPnet’s result (E) and RELION’s result (F), respectively. The particles’ angular distributions in reconstruction process are displayed in Rot-Tilt angle scatter plots by results of DRPnet (G) and RELION (H). A gray “X” symbol indicates a missing view. The last sub-figure overlays the histogram of Max Value Probability Distribution (I) of particles picked by DRPnet (blue dashed line) and RELION (red solid line).

**Figure 6:**
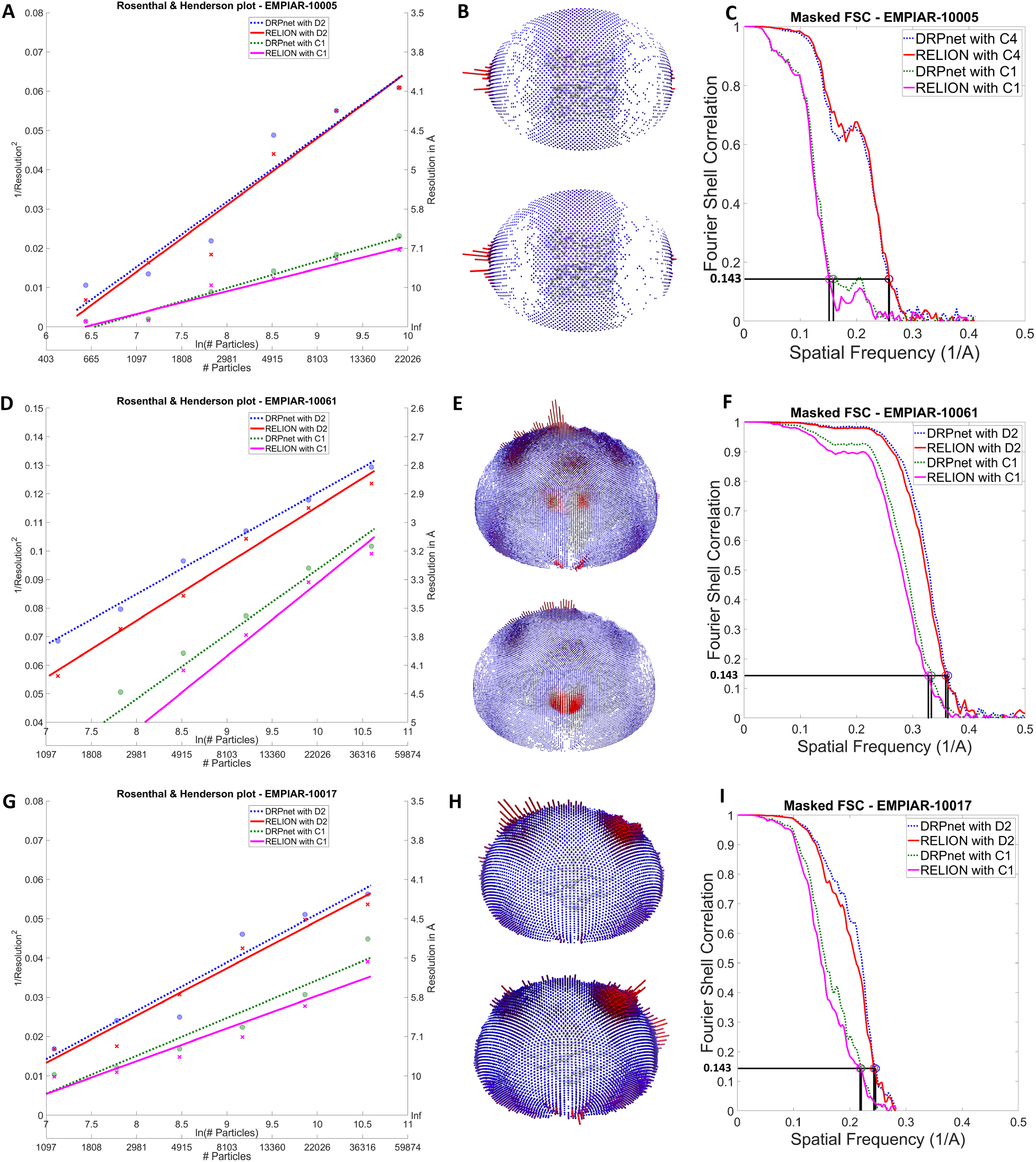
3D reconstruction outcomes: the right column contains Fourier Shell Correlation curves, the middle column contains visualization of bild files for the angle distributions of good particles picked from 3 datasets by DRPnet’s pick (upper) and RELION’s autopick (lower), and the left column contains Rosenthal & Henderson plots show how resolution of 3D structures changed when the number of picked particles changed. The first row shows EMPIAR-10005 (TRPV1)’s results: (A) Rosenthal & Henderson plot, (B) angle distribution by DRPnet’s pick (upper) and RELION (lower), (C) Masked FCS curve; The second has EMPIAR-10061 (*β*-galactosidase) results: (C) Rosenthal & Henderson plot, (D) angle distribution by DRPnet’s pick (upper) and RELION (lower), (F) Masked FCS curve; The last row shows EMPIAR-10017 (*β*-galactosidase)’s results: (G) Rosenthal & Henderson plot, (H) angle distribution by DRPnet’s pick (upper) and RELION (lower), (I) Masked FCS curve.

## 5 Conclusions

In summary, we propose a Deep Regression Picker Network (DRPnet) and successfully demonstrate the ability of picking particles on a dataset of cryo-EM micrographs which is different from the training data. Our deep learning based network is simple and effective for automatically picking particles from 2D cryo-EM micrographs, including those with low contrast or having a large particle box size.

DRPnet has a two-part cascade of convolutional neural networks for detection and classification. It provides flexibility to retrain either or both networks if desired. Compared to other deep learning tools for particle picking in cryoEM micrographs reported to date, this cascade strategy is unique. After the initial training with TRPV1, we found it was unnecessary to retrain this network when applied to other cryoEM micrographs containing differently shaped particles (e.g. *β*-galactosidase) in other datasets (EMPIAR-10061). Without the need to retrain, automating 2D template preparation to reconstruct 3D protein structures by single particle analysis is feasible. If desired, one can retrain the classification CNN – which may be useful for very low contrast images or images that have artifacts, such as shapes similar to particles of interest.

Additionally, utilizing a pretrained network to pick particles significantly reduces time to obtain a particle dataset. DRPnet is very efficient for selecting a set of particles. For comparison, picking ~1000 particles manually in our laboratory from the above EMPIAR datasets took about 30 minutes per dataset. There is also additional time time spent running 2D classification to select good 2D class average templates, as well as optimizing autopicking parameters. With DRPnet, it takes about 30 seconds per micrograph to pick the particles, which results in significant time savings.

When investigating traditional computer science identification metrics of recall, precision, and F-measure, we did not feel these indicators were informative of performance. Many times, existing ground truth sets are lacking due to 1) low contrast particles not contained in the “experts” ground truth, or 2) the particles composing final 3D reconstruction not representing the total particles used for alignment, classification, and averaging. Specifically, in case of EMPIAR-10017 (*β*-galactosidase), although the ground truth was the most comprehensive, it still did not cover all particle patterns, thus orientations. This may be due to fluctuations in intensity level, which may effect the ground truth’s particle patterns. Cryo-electron microscopy images captured with low contrast (low defocus or minimal electron radiation exposure, therefore enhancing high resolution terms) are most likely have high resolution 3D protein structural information, and conversely, those captured with high contrast (high defocus or high electron radiation exposure, enhancing low resolution terms) contain low resolution inforation. Because images in EMPIAR-10017 (*β*-galactosidase) have different contrast levels, a true positive particle in a low contrast image can be a false positive pattern in high contrast image. For the above reasons, the detection stage F-measure of 78.54% was not high, but it was the best result we could achieve with EMPIAR-10017 (*β*-galactosidase) data.

Therefore, we sought additional metrics to guide our analysis. Metrics we found useful include reporting resolution using the gold standard Fourier Shell Correlation (correlation threshold of 0.143) [50], angular distributions plots, efficiency (E_od_) value [49], plots of maximum value of probability distributions, and inspecting the 3D isosurface.

Gold-standard Fourier Shell Correlation or 3D reconstruction resolution reveals the internal consistency in 3D reconstruction output maps [50]. Consistently, we obtained slightly higher resolutions with DRPnet over Relion’s autopick, and DRPnet excelled with low contrast data sets (those collected on Gatan direct detectors with low defocus values). Plotting the angular distributions allows one to visually inspect the orientations (angles and rotations) of the picked particles and also their abundance. With DRPnet, there is more coverage (less white background) in all datasets. This finding is also corroborated by the efficiency metric, with DRPnet consistently showing improved E_od_ values, indicating a more robust sampling of particle angular orientations in the final 3D reconstruction. When assigning a particle’s orientation during 3D refinement, histogram plots of the Maximum Probability Distribution Values provide a reference of how confidently a particle’s orientation was assigned (a value of 1 indicates that the probability value has converged, and a value of 0 indicates great uncertainty) [64]. The shape of this distribution is informative; we note that with DRPnet-picked particles for EMPIAR-10061 & EMPIAR-10005, there is a greater number of particles having a higher maximum probability distribution value, with the shape of the histogram distribution shifted to the right. For our DRPnet analysis of EMPIAR-100017, the two curves are very similar; with RELION’s autopicked particle distribution values having slight improvement over DRPnet’s. Finally, visual inspection of the 3D reconstruction isosurface with USCF Chimera [65] or generating difference densities revealed little to no improvement in secondary structure and side chain densities with DRPnet versus RELION autopicked 3D reconstructions. 3

We noted minor improvements in 3D reconstruction resolution with DRPnet as compared to RELION’s picking strategy, and believe this is due to picked particles having better angular orientation coverage. This was true for both TRPV1 (EMPIAR-10005) and *β*-galactosidase (EMPIAR-10061), which are low contrast (low defocus, high resolution) datasets collected from Gatan direct detectors. Based on the concept of blob detection, DRPnet can pick diversified patterns which usually results in wider particle orientation angle coverage, allowing an improvement in resolution. This is different than the manually selected templates used in RELION’s autopicking function. It is likely especially for low contrast (low defocus, high resolution) datasets that bias from the user is minimized, preferred high contrast particle views are picked as well as other less obvious views with less contrast with the resulting particle set having more diverse angular orientations. We also note that in these aforementioned results, we used a subset of the total data. When using an entire EMPIAR-10017 (*β*-galactosidase) dataset to test, containing numerous particles (around 50,000) to perform 3D reconstructions, DRPnet and RELION reached the same resolution. If limited by low contrast data or not enough particles, DRPnet may be particularly useful.

Here, we present deep regression-based particle picking algorithms composed of a dual cascade of convolutional neural networks, entitled “DRPnet”. The power of this strategy is that 1) the picking algorithm does not have to be, but can be, retrained, 2) efficiency of automated particle picking compared to manual template generation, and 3) usefulness for low contrast datasets (low defocus, high resolution) or data that is not complete (needs better angular sampling). We also provide functional assessment metrics for cryoEM micrograph particle picking beyond traditional recall, precision, and F-measure. In the future, we anticipate developing a powerful classification network for our workflow, trained with multiple types of particles (heterogeneous populations of particles, for example). Having a more powerful classification would also reduce the running time of our picking program while increasing its accuracy (reducing false positive particles such as noisy and/or low contrast particles). The second issue we also need to improve is more accurate localization of the particles within the image patches. Tackling these issues will improve the DRPnet algorithm and also will provide a refined dataset to perform high resolution 3D reconstructions.

## Notes

#### Summary of Updates

Change color maps from Jet to Parula, and change some minor details.

